# Notch controls early temporal factor expression to control timing of Mushroom body neuroblast apoptosis

**DOI:** 10.1101/2024.01.31.578279

**Authors:** Kendall R. Branham, Chhavi Sood, Susan E. Doyle, Matt C. Pahl, Sarah E. Siegrist

## Abstract

The neurogenic period, where neural stem cells (NSCs) proliferate to produce molecularly distinct progeny in the developing brain, is a critical time of growth in many organisms. Proper brain development is crucial for survival and requires strict regulation of NSC divisions along a set developmental timeline. In *Drosophila* NSCs known as neuroblasts (NBs), cell intrinsic programs integrate with extrinsic cues to control periods of rapid growth through temporal patterning genes. Without regulation, NSCs can under proliferate leading to diseases like microcephaly and autism spectrum disorders, or over proliferate leading to macrocephaly and tumors. We know programs to control timing of proliferation and elimination of NSCs exist, but many elements of temporal cassettes are still unclear. What genes may be upstream to regulate known temporal programs to control when certain progeny are produced have not been fully identified, leaving a gap in our understanding. To address these questions, we carried out a large-scale RNAi screen aimed at identifying genes required for NSC elimination. We identified Notch and its ligand, Delta. When Notch pathway activity is reduced in NSCs, we found premature elimination of an important subset of neuroblasts called the mushroom body neuroblasts (MBNBs). These MBNBs produce the neurons responsible for formation of the evolutionarily conserved structure called the mushroom body (MB), which is involved in olfactory based learning and memory. MBNBs with reduced Notch pathway activity also experienced defects in MB structure. Furthermore, we determined that temporal patterning is disrupted primarily through loss of early temporal factor expression. In this work, we find that cell signaling pathways that involve the receptor Notch and its ligand. Delta function to regulate NB proliferation in *Drosophila melanogaster* by regulating early temporal factor expression.

## Background

Stem cell division during development is a fundamental biological process that occurs across all organisms and requires strict regulation for appropriate tissue construction. In the developing central nervous system (CNS), proper timing of neural stem cell (NSC) proliferation during development is paramount to the formation of morphologically correct and functional neural tissues (Hartenstein & Wodarz, 2013; Matsubara et al., 2021). If NSC divisions are not regulated, under proliferation can occur leading to reduced neuron numbers and microcephaly (Courchesne et al., 2018; Gilmore & Walsh, 2013). Conversely, over proliferation of NSCs can result in excess growth leading to macrocephaly. Both scenarios of under and over proliferation can lead to neurodevelopmental disorders, including autism spectrum disorders (ASD) (Parenti et al., 2020; Katsimpardi & Lledo, 2018; Courchesne et al., 2018). Therefore, initiating divisions as well as terminating divisions of NSCs at proper points in development are both equally important for brain and neural tissue formation.

It is understood that cell intrinsic factors integrate their signaling with extrinsic cues to regulate NSC proliferation. Timed expression of specific temporal patterning genes allows for production of molecularly distinct progeny through development to form all the cell types needed for a functional brain. Transient activation of temporal patterning genes is controlled by temporal cassettes consisting of temporal transcription factors (tTFs). These tTFs respond to extrinsic cues such as steroid hormones and surrounding glial cell signals (Liu et al., 2019; Rossi & Desplan, 2020; Syed et al., 2017; Sood et al., 2023). It is crucial to understand how systems of intrinsic and extrinsic cell signaling interact to regulate gene expression and ensure correct proliferation and differentiation of NSCs in the developing brain, as well as how they initiate termination of NSCs when development is complete.

The regulation of neurogenesis through temporal patterning is seen across species, including in the fruit fly *Drosophila melanogaster*. The neural development of the fruit fly brain is tightly regulated to correctly pattern the growing brain throughout its different life stages. NSCs in *D. melanogaster* are known as neuroblasts (NBs) and have distinct subtypes that produce the varying progeny needed for the adult fly brain (Bayraktar & Doe, 2013; Doe, 2017; Islam & Erclik, 2022; Rossi et al., 2017). A particular neuroblast subset of interest within the central brain are the mushroom body neuroblasts (MBNBs). MBNBs give rise to the evolutionarily conserved mushroom body (MB) structure known to function in learning and memory, particularly when integrated with olfaction (Kunz et al., 2012; Kurusu et al., 2000; Lee et al., 2020, Lin, 2023). The four MBNBs and of each fly central brain hemisphere actively proliferate throughout development, from embryonic stages until late pupal stages (Ito & Hotta, 1992; Kunz et al., 2012). This differs from other central brain neuroblasts (CBNBs) that enter a period of quiescence during early larval stages, reactivate in response to nutritional cues to divide again, then permanently terminate their divisions during early pupal stages (Hartenstein & Wodarz, 2013; Nassel et al., 2015). MBNBs are considered type I neuroblasts which means they asymmetrically divide to both self-renew and produce an intermediate progenitor called a ganglion mother cell (GMC). GMCs then symmetrically divide to produce neurons, called Kenyon cells (KCs), that are sequentially produced, forming the distinct γ (early born), ⍺’/β’ (middle born), and ⍺/β (late born) neurons of the mushroom body (Ito et al., 1997; Liu et al., 2015). KCs extend their dendrites to an area on the dorsal surface called the calyx, and their axons traverse the brain through the peduncle and bifurcate to form the different lobes. Intrinsic temporal programs determine what type of KC is produced when in response to changes in temporal programs triggered by extrinsic signaling cues (Heisenberg, 2003; Rossi & Desplan, 2020). The final MB is a five-lobed structure consisting of ∼2,000 neurons produced from the four MBNBs of each hemisphere. After forming this structure, MBNBs are eliminated via parallel pathways of both apoptosis and autophagy prior to adulthood (Siegrist et al., 2010; Pahl et al., 2019).

Temporal patterning genes like hunchback (Hb), kruppel (Kr), pdm, castor (cas), and grainy head (Grh) have been shown to regulate periods of embryonic neurogenesis and types of neurons produced, but outside of embryonic stages temporal patterning genes and their programs are less understood (Brody & Odenwald, 2000; Isshiki et al., 2001; Li et al., 2013; Kohwi & Doe, 2013). It is currently known that an opposing gradient of RNA binding proteins Insulin-like growth factor II (Imp) and Syncrip (Syp) are expressed in MBNBs and are responsible for patterning MBNB progeny (Liu et al., 2015; Rossi et al., 2021). During early developmental stages when Imp is expressed, chronologically inappropriate morphogenesis (Chinmo) and Lin-28 positive γ neurons are produced, whereas in later stages when Syp is expressed, Let-7 and Broad (Br) positive progeny are produced (Doe, 2006; Islam & Erclik, 2022). Chinmo is a BTB zinc-finger (BTB-ZF) transcription factor whose levels are high in early born γ neurons but decrease to no expression in late born ⍺/β neurons due in part to activity of the microRNA Let-7 (Zhu et al., 2006; Wu et al., 2012; Liu et al., 2015; Islam & Erclik, 2022). As levels of early factor expression decrease in MBNBs, late factor expression increases. Syp represses early factor expression and promotes even later temporal factor expression and during the overlap of temporal factors (low Imp and increasing Syp). During this period of temporal factor overlap, ⍺’/β’ neurons are produced that are maternal gene require for meiosis (Mamo) positive (Rossi & Desplan, 2020). Mamo, is a BTB-ZF transcription factor whose expression is stabilized by the late factor Syp (Liu et al., 2015; Rossi & Desplan, 2020). Finally, expression of Ecdysone induced protein 93 (E93) is positively regulated by Syp and ecdysone steroid hormone signaling. E93 inhibits MBNB PI3-kinase activity to activate autophagy and prime MBNBs for elimination via apoptosis (Siegrist et al., 2010; Pahl et al., 2019).

While it is clear that Imp and Syp and their targets function to determine types of progenies produced over time, it still remains unclear how forward progression through temporal programs are controlled. To better understand regulation of temporal patterning, we carried out a targeted RNAi screen aimed at identifying genes required to terminate MB neurogenesis on time. From this screen, we identified the transmembrane receptor, Notch and its ligand, Delta. Notch binds to its membrane bound ligand Delta expressed on neighboring cells. After ligand binding, Notch is proteolytically cleaved allowing for the nuclear translocation of the Notch intracellular domain (NICD) to the nucleus to regulate gene expression (Schnute et al., 2018; Kandachar & Roegiers, 2012; van Teetering et al., 2009). Notch signaling is evolutionarily conserved and during development, has known roles in regulating binary cell fate choices (Sato et al., 2016). Notch also controls cell cycle exit to promote NB quiescence during the embryonic to larval transition and regulates termination of neurogenesis of CBNBs outside the MBNBs (Sood et al., 2022; Sood et al., 2023). Here we report the role of Notch and its associated ligand Delta in temporal patterning and timing of MB neurogenesis termination.

## Results

### Notch Pathway Knockdown Results in Premature Loss of MBNBs

MBNBS are known to actively proliferate from embryogenesis until 84 to 90 hours after pupal formation (APF), shortly before adult eclosion. To understand the role that Notch pathway signaling plays in maintenance of this division period, we knocked down Notch (N) and its ligand Delta (Dl) in all neuroblasts by driving expression of *UAS-NRNAi* or *UAS-DlRNAi* with a neuroblast specific driver *worGal4* (hereafter referred to as *NB Gal4*) and examined MBNB number during pupal phases. At 48 h APF, no difference was observed in the number of MBNBs between *NRNAi* and control animals (Fig. 1A-B, G, MBNBs marked with arrowhead). However, a significant reduction of MBNBs was observed following Delta knockdown (Fig. 1C, 1G). By 72 h APF we found a significant reduction of MBNBs in both Notch and Delta knockdown brains compared to control animals (*NRNAi*, 3.7 ± .13, *DlRNAi*, 2.1 ± .31 MBNBs compared to controls, 4 ± .19 MBNBs) (Fig. 1D-F, G). Note the presence of other CBNBs in *DlRNAi* animals (see discussion). Furthermore, cell size based on the average diameter was reduced in MBNBs still present in *NRNAi* and *DlRNAi* brains compared to controls. At 48 h APF, all genotypes tested showed MBNBs with an average diameter size of 10 μm. However, by 72 h APF, *NRNAi* brains had MBNBs with an average diameter of 8.5 ± .14 μm and *DlRNAi* brains with an average of 7.0 ± .27 μm. (Fig. 1H). The observation of significantly smaller sized MBNBs in *NRNAi* and *DlRNAi* animals suggests that these MBNBs still undergo normal reductive cell size divisions before their elimination. Reductive cell division occurs in wild type animals in part through autophagy in preparation for apoptotic cell death as levels of PI3K decrease. Indeed, at 72 h APF in *DlRNAi* brains, we observed morphological characteristics of apoptotic cell death in MBNBs (Fig 1F inset). To further confirm these results, Notch loss of function allele *Notch55e11* MARCM clones were employed to assess how a null allele for Notch would affect MBNB survival. At 48 h APF we found that the MBNB in the Notch55e11 MARCM clone was absent while control MBNBs were still present (Supp. 1). We conclude that the Notch pathway signaling regulates timing of MB neurogenesis, and when Notch pathway activity is attenuated, MBNBs are prematurely lost, possibly through apoptosis.

**Figure 1.**
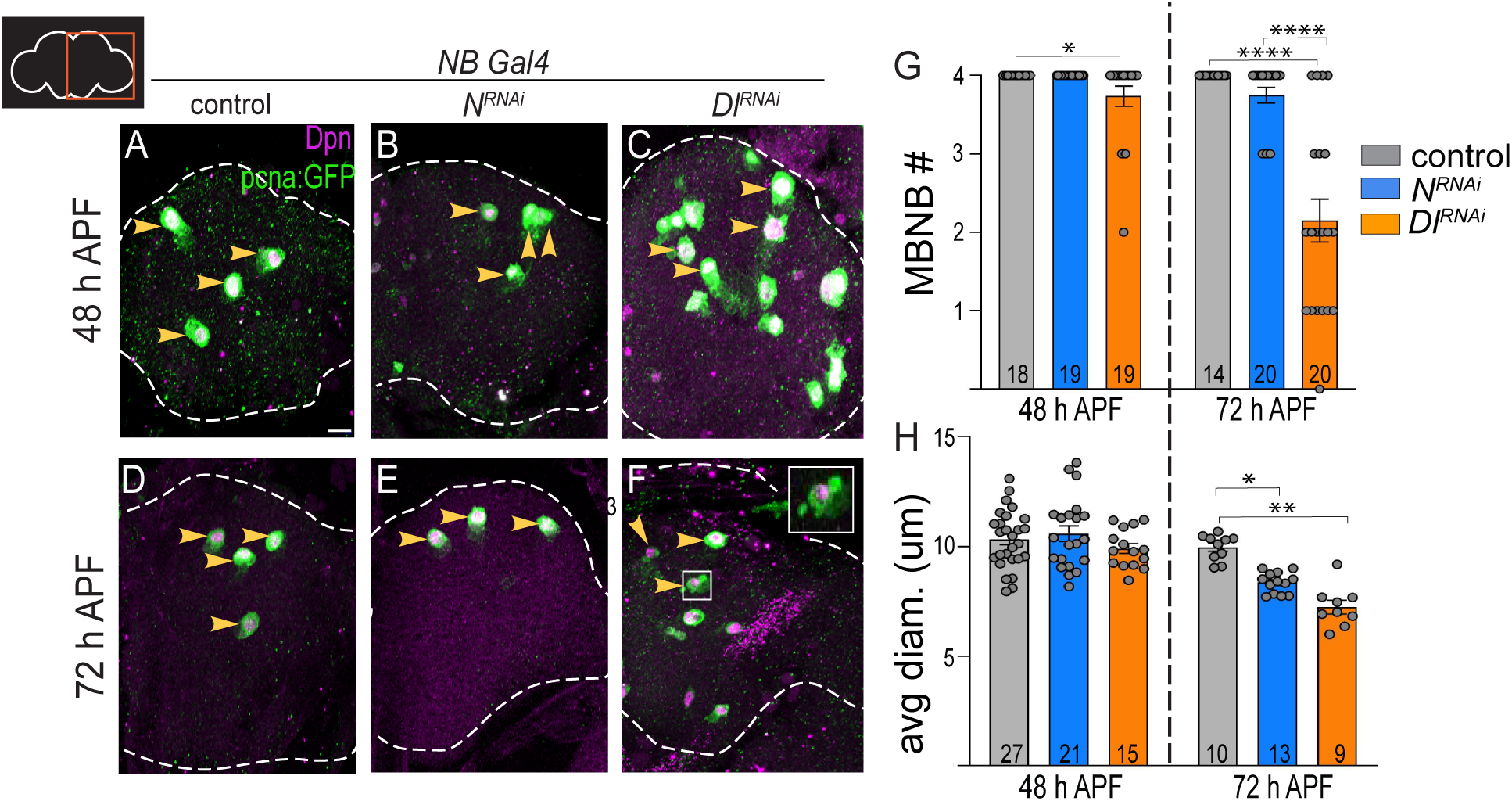
Notch pathway knockdown results in premature loss of MBNBs. (A-F) Cartoon in top left indicates brain hemisphere imaged for this and all subsequent figures. Maximum intensity projections of the right brain hemisphere with pupal timepoints and genotypes as listed. Antibodies used are indicated in the top right of panel A and the hemisphere is outlined in a white dashed line. Yellow arrows indicate MBNBs, unmarked NBs are other non-MBNB central brain NBs. The inset (F) indicates a MBNB undergoing premature cell death. Scale bar = 20 μm. (G) Histogram represents the mean number of MBNBs present at the stated time points for each genotype. Each data point represents a separate brain hemisphere with the n represented within the bar. (H) Histogram represents the average MBNB diameter at the stated time points for each genotype. Each data point represents an individual MBNB with the n represented within the bar. (G-H) Error bars represent the SEM, alpha = 0.05, analyzed with a one-way ANOVA followed by a test for multiple comparisons. Significance is indicated with asterisks.

### *DlRNAi* MBNBs undergo premature cell death

Because observed phenotypes of early termination and reduced cell size were more penetrant in *DlRNAi* lines than *NRNAi*, we chose to focus on *DlRNAi* animals for the subsequent analyses. In wild type animals, MBNBs are eliminated via autophagy/apoptosis during late pupal stages whereas other CBNBs terminally differentiate during early pupal stages (Siegrist et al., 2010; Pahl et al., 2019; Maurange et al., 2008). When Notch pathway activity is reduced, MBNBs are lost prematurely with some displaying blebbing cell morphology consistent with apoptosis. To determine whether *DlRNAi* MBNBs are eliminated via cell death as in control animals, or change their mechanism to terminal differentiation, we co-expressed an inhibitor of apoptosis (*UAS-miRHG)* in *DlRNAi* animals. In one day old *DlRNAi* adults, no MBNBs are observed (Fig. 2A, C). Yet in *DlRNAi* animals that co-express an inhibitor of cell death, we observed a significant number of MBNBs (Fig. 2B, C). On average, 2.5 ± .29 MBNBs were seen in one day old adults compared to 0 MBNBs in brains with *DlRNAi* alone (Fig. 2C). We conclude that MBNBs undergo apoptosis when Notch pathway activity is reduced, however MBNB elimination via apoptosis occurs prematurely.

**Figure 2.**
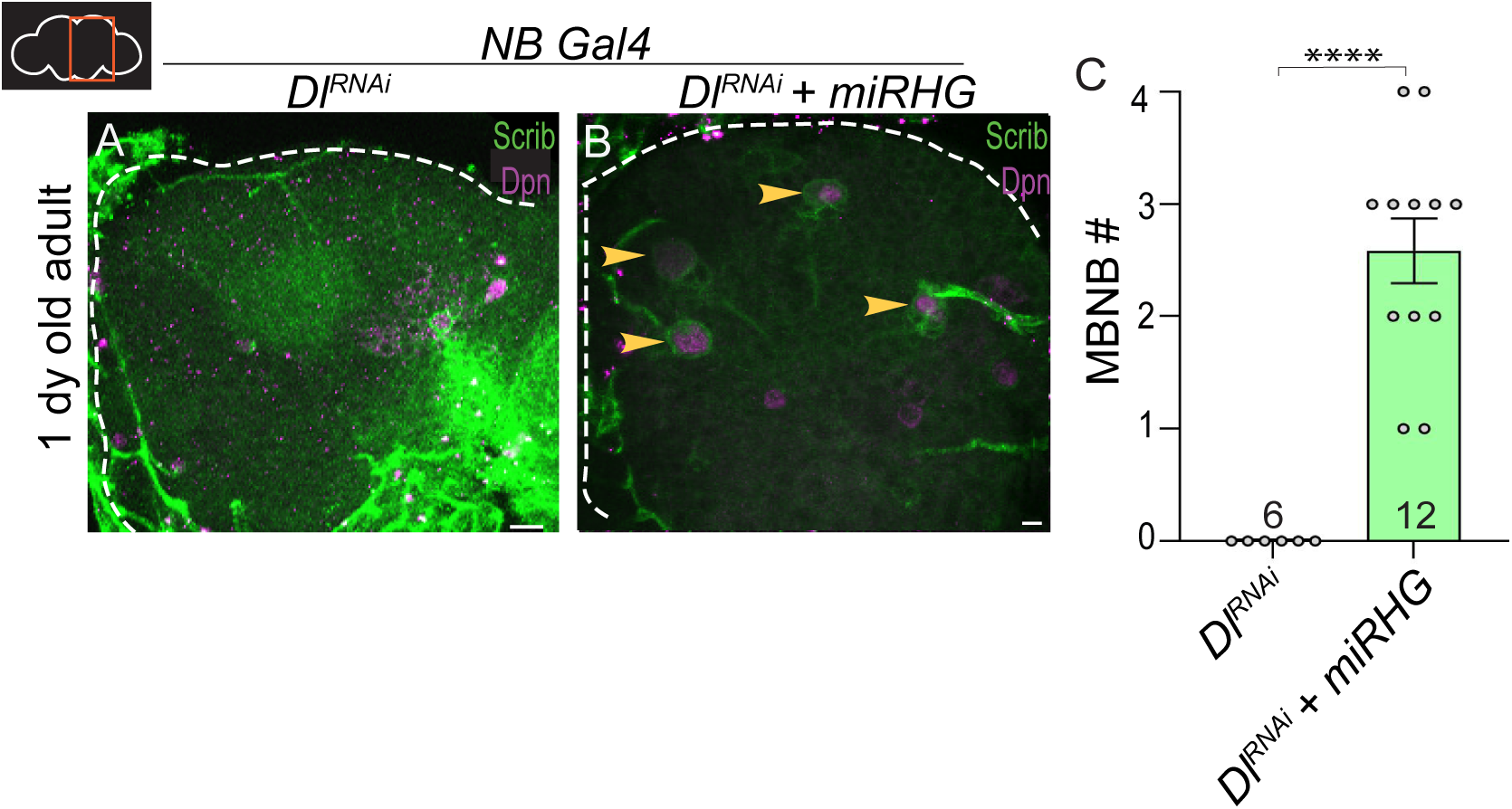
Blocking apoptotic cell death in a *DeltaRNAi* animal allows long term persistence of MBNBs. (A) Maximum intensity projection of the right hemisphere of 1 day old adult brains of the specified genotypes. Scale bar = 10 μm. (B) Maximum intensity projection of the right hemisphere of freshly eclosed brains of the specified genotype. Persisting MBNBs indicated with yellow arrowheads. Scale bar = 10m. Markers used are indicated under the corresponding panels. (C) Histogram representing the average number of persisting MBNBs at the FE time point in *DeltaRNAi* and *DeltaRNAi + UAS-miRHG* brains. Each data point represents one brain hemisphere for a total of 12 brain hemispheres from 6 different animals. Data was analyzed using an unpaired, two-tailed student’s t-test, alpha = 0.05. Error bars represent the standard error of the mean. Significance is indicated with asterisks.

### Temporal Windows are Shifted in Notch Pathway Knockdown

MBNBs as opposed to other CBNBs are known to experience a longer period of Imp expression (Liu et al., 2015). This prolonged Imp expression allows for extended proliferation, accounting for the longer life cycle of MBNBs compared to other CBNBs. In NBs, the Imp to Syp transition is mediated by the steroid hormone, ecdysone (Doe, 2017). However, it remains unclear whether other factors are required. After observing premature loss of MBNBs in both *NRNAi* and *DlRNAi*, we hypothesized that temporal patterning may be altered given that MBNBs are also lost prematurely when Imp is reduced (Liu et al., 2015). To test this hypothesis, we employed heat shock flippase clone constructs which allows for expression of *DlRNAi* and a fluorescent protein in a subset of MBNBs. This technique provides a control wild type MBNB alongside a *DlRNAi*, GFP expressing MBNB clone for direct comparison within the same brain hemisphere. Brains were staged upon hatching, heat shocked, and then imaged at 72 h after larval hatching (ALH).

Imp expression in *DlRNAi* clones was reduced compared to wildtype MBNBs (Fig. 3A, yellow arrowhead). Approximately 54% of MBNBs expressing *DlRNA*i maintained Imp expression compared to 93% of control MBNBs (Fig. 3E). This result indicates that Notch/Delta signaling positively regulates Imp expression. Because Imp and Syp reciprocally inhibit one another (as Imp decreases, Syp increases), we tested whether premature loss of Imp would lead to precocious Syp expression (Liu et al., 2015). We again used *DlRNAi* clones to assess Syp expression at 72 h ALH. As expected, no Syp expression was seen in controls as the Imp window is still active at this time (Fig. 3B, red arrowhead). However, precocious Syp expression was also not observed in MBNB *DlRNAi* clones (Fig. 3B, yellow arrowhead, n = 6). We do see Syp expression in whole brain knockdown of Dl at 48 h APF, leading to the assumption that syp expression may just be delayed without Dl (Fig 3D). We conclude that Notch/Delta signaling positively regulates early temporal patterning by promoting Imp expression. Moreover, additional cues are likely required to promote Syp, cues that are independent of early temporal factor Imp expression.

**Figure 3.**
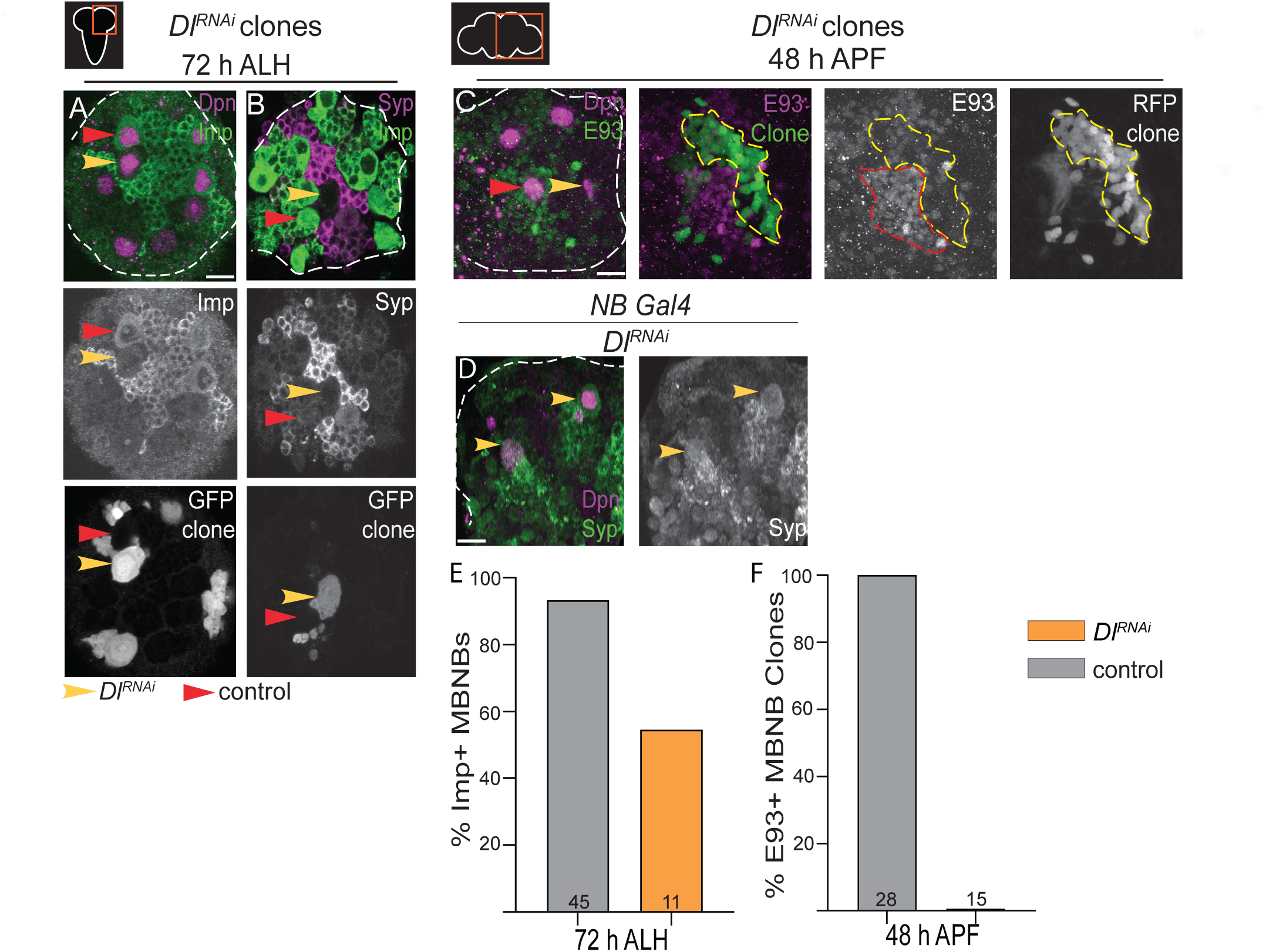
Delta knockdown alters temporal factor expression in MBNBs. (A-C) Maximum intensity projections of the right brain hemisphere with larval and pupal timepoints and genotypes as listed. Markers are indicated in the top right of the corresponding panels. Single channel grayscale images provided to show clone MBNB and temporal factor being analyzed. Control MBNBs are indicated with red arrowheads while clones are indicated with yellow arrowheads. Brain hemispheres are outlined with white dashed lines. The Delta clone stained for E93 is outlined in a yellow dashed line. The control MBNB’s E93 is outlined in a red dashed line. Scale bar = 10μm. (D-E) Histogram showing the percentage of imp and E93 positive control MBNBs versus clone MBNBs or progeny. A count of syp positive cells is not shown as it was zero for both classes of MBNBs. The number of each type of MBNB quantified is listed within the bar. The value was calculated by dividing the number of positive cells by the total number of cells present at the particular time point.

Since early temporal patterning is altered in *DlRNAi* MB NB clones, we next asked whether late temporal patterning is also altered. At 48 h APF, control MBNBs express low levels of E93 but high levels of E93 in their Kenyon cell progeny (Fig. 3C, red arrowhead and C’, red outline). However, at 48 h APF in *DlRNAi* clones, E93 was not detected in MBNBs nor in their Kenyon cell progeny (Fig. 3C, yellow arrowhead and 3’, yellow outline). This is drastically different from the 100% of control MBNBs expressing E93 at this time point (Fig. 3F). Previous work has shown that expression of both the intrinsic temporal factor Syp and the extrinsic cue ecdysone is important for promoting E93 expression in MBNBs at late developmental stages prior to termination (Pahl et al., 2019).

### Mushroom Body Defects are seen in Notch Pathway Knockdown

Proper mushroom body morphology is dependent on MBNBs proliferating and producing the correct subtype of neuronal progeny at distinct timepoints of the neurogenic period. We found that when MBNB proliferation is disrupted by altering Notch pathway signaling, the MB structure is severely affected. We stained freshly eclosed adult brains with the MB marker Fasciclin 2 (FasII) that labels the γ and ⍺/β neurons and a membrane marker Scribble (scrib). Compared to controls, we see in *DlRNAi* animals that late born ⍺ and β neurons are absent and early born γ neurons make up a large portion of the structure (Fig. 4A-D). While γ cells are present, the γ lobe does not look like it has undergone the same amount of axonal pruning as would be expected at the adult stage (Fig. 4D-D’’) (Yu & Schuldiner, 2014). These abnormal looking structures appear to result from defects in γ neuron differentiation based on morphology (Figure 4C, yellow arrowhead). MB neurons are experiencing defects in molecular identity and number. The survival of these neurons is affected by loss of delta which would account for absence of late born kenyon cell types. Observed morphological defects show the drastic effect Notch pathway disruption can have on important structures in the adult brain.

**Figure 4.**
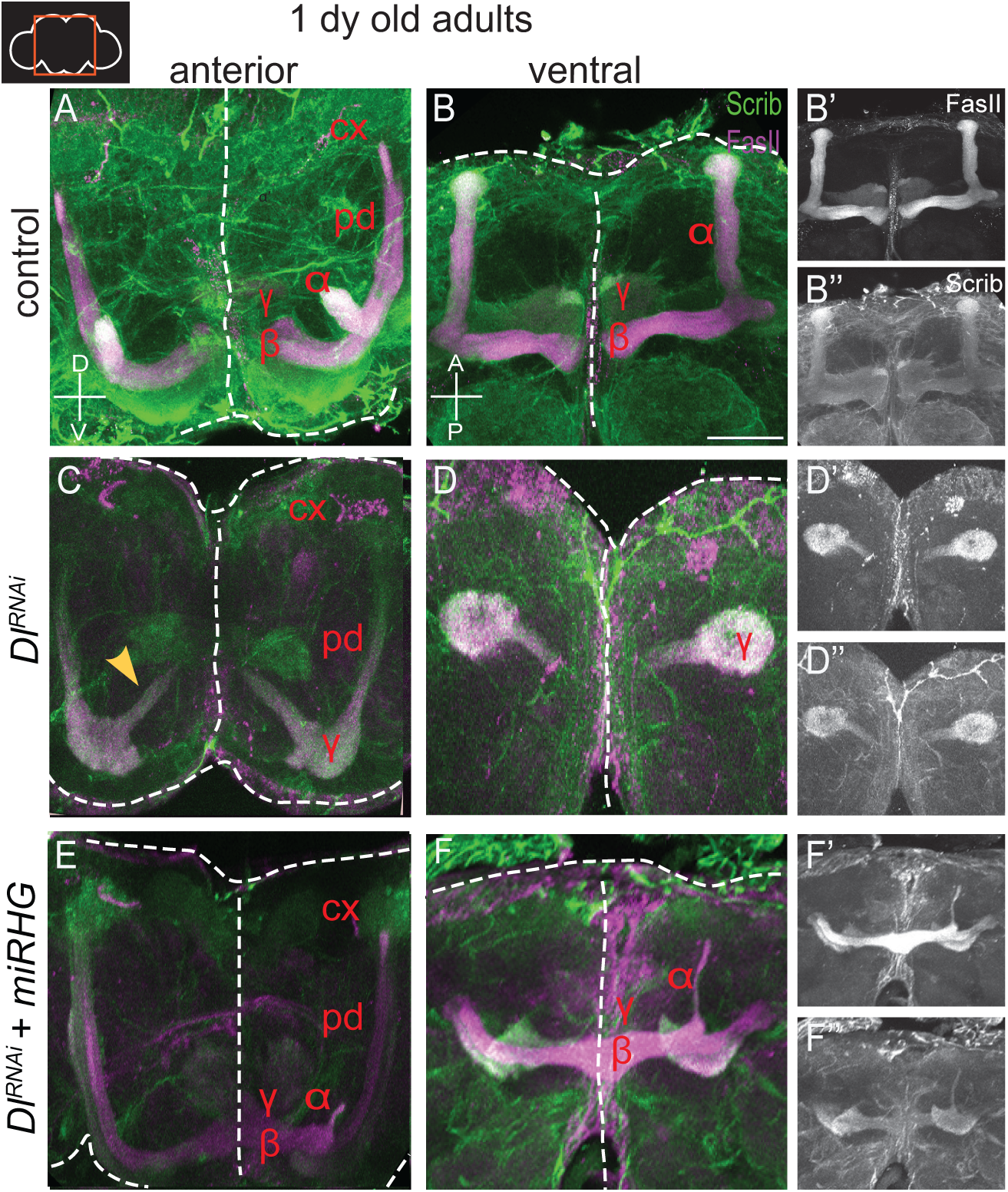
Premature loss of MBNBs causes severe MB morphological defects. (A-F) Maximum intensity projections of the central brain lobes of one day old adult brains of the specified genotypes. White dashed lines outline the central brain and indicate the midline. Images show expression of alpha, beta, gamma neuron marker fasII and scrib. The ⍺’ and β’ lobes are not labeled. Red lettering indicates which lobe is present for each genotype. (A, C, E) Anterior view lobes are marked to indicate the alpha, beta, gamma lobes as well as the peduncle and calyx when applicable. (B, D, F) Ventral view lobes are marked to indicate the alpha, beta, gamma lobes. (B’-F’’) Single channel grayscale images show the MB structure from the ventral view for both FasII and Scrib expression. Scale bar = 50 μm. (A-B) n= 12. (C-D) n=4. (E-F) n = 3.

Following our results found in figure 2, we wanted to test if blocking death in *DeltaRNAi* animals by using the *UAS-miRHG* construct would restore some of the MB structure in adult animals. We found that when apoptosis was inhibited, the MB morphology is not identical to control, but appears to have better organization of the γ neurons than in *DeltaRNAi* brains (Fig. 4E-F’’). In addition, we see a slight increase in the production of late born, ⍺ and β neurons, enough to form an ⍺ lobe to some extent (Fig. 4F). While we did not observe a complete rescue of MB morphology, this finding further emphasizes that when death is blocked in a Delta knockdown animal, MBNBs are able to persist longer. Their lineage is then able to better survive and can integrate into the MB structure.

### Eyeless as an Upstream Regulator of Notch

An open question that remains is how Notch differentially controls NB lineage progression and timing of neurogenesis termination. In MB NBs, when Notch activity is reduced Imp expression is lost prematurely, leading to premature neurogenesis termination, whereas in the other CBNBs (∼100), Imp expression is prolonged and neurogenesis is extended (Sood et al., 2023). To understand these lineage specific differences in response to Notch activity, we asked whether Eyeless is required. Ey is a paired box homeodomain transcription factor known for its role in eye development (Martini et al., 2000; Noveen et al., 2000). It is highly conserved and known as Pax6 in mammals (Callearts et al., 2001). To assess the role of Ey in mediating Notch activity in MBNBs we used the heat-shock flippase induced clone system again with *UAS-EyRNAi* and a Notch activity reporter, *enhancer of split (E(spl))-GFP.* At 24 h APF and 72 h APF, *EyRNAi* clones had a reduction in fluorescence from the E(spl)GFP reporter, suggesting that Notch activity is reduced compared to control MBNBs (Fig 5A-B). We calculated the percent of MBNBs expressing E(spl)GFP at 72 h APF and found that only 40% of *EyRNAi* clones are positive for GFP while 100% of control MBNBs are positive for GFP (Fig 5C). This supports the possibility that Ey may regulate Notch activity and may be a key gene in establishing lineage specific differences between MBNBs and other CBNBs.

**Figure 5.**
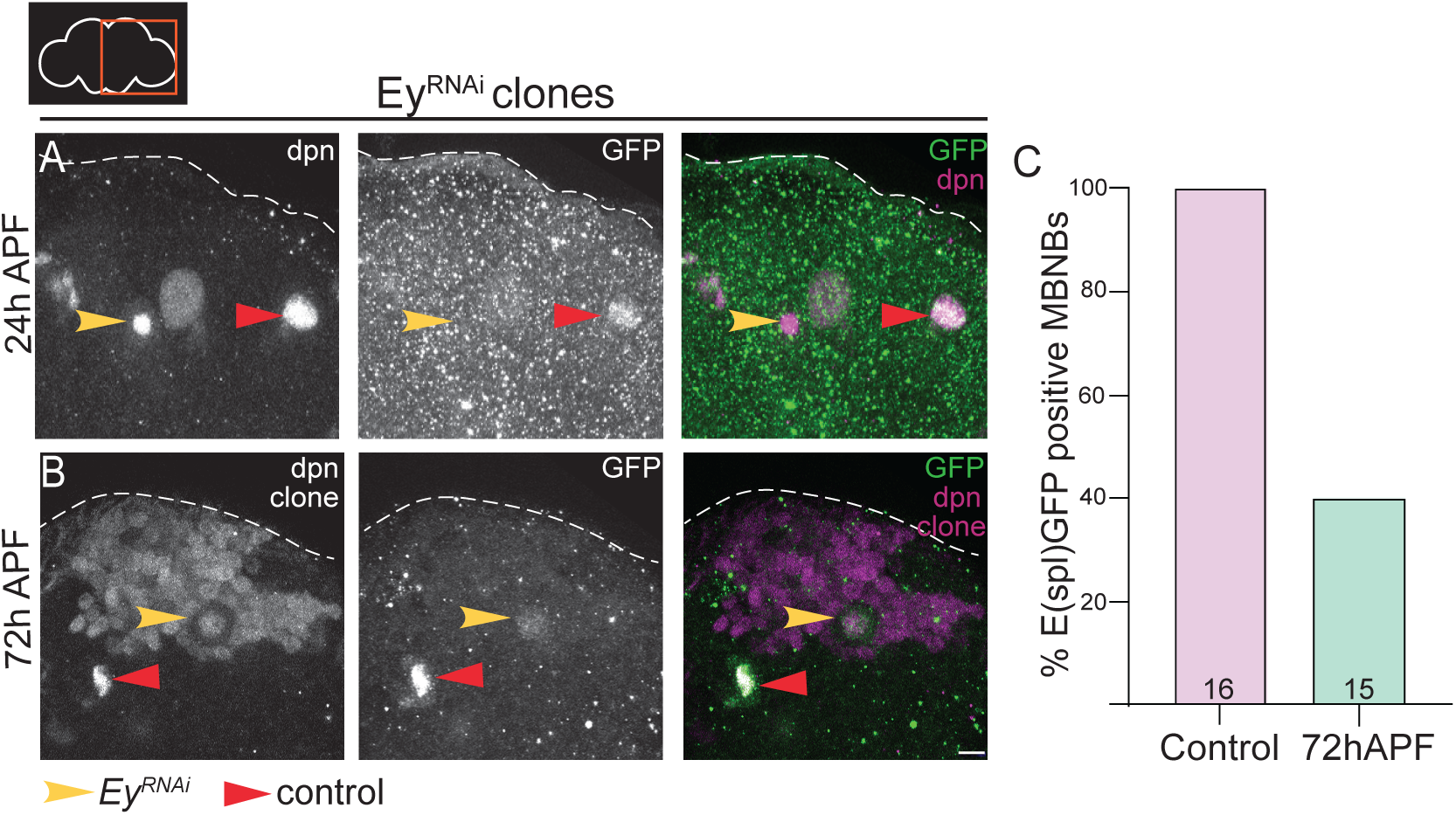
Eyeless regulates Notch activity in MBNBs. (A-B) Single z-stack images of the right brain hemisphere at time points as indicated to the left of images. *EyRNAi;E(spl)GFP* MBNBs clones are indicated with a yellow arrowhead while control MBNBs are indicated with a red arrowhead. Brain hemispheres are outlined with white dashed lines. Scale bar = 10 μm. Markers used are indicated under the corresponding panels. (C) Histogram showing the percentage of *E(spl)GFP* positive control MBNBs versus clone MBNBs. The amount of each type of MBNB quantified is listed within the bar. The value was calculated by dividing the number of positive cells by the total number of cells present at the particular time point. A total of 9 hemispheres from 5 different animals was quantified.

## Discussion

### Notch activity is lineage-specific

Here we report that Notch pathway signaling regulates the proliferative window of MBNBs in the developing *D. melanogaster* brain. We found that when Notch pathway components are knocked down using RNAi lines against Notch receptor and its ligand Delta, MBNBs prematurely terminate, and the MB structure is disrupted. This is due to a shift in the known Imp-Syp-E93 temporal cassette. Rather than having a long period of Imp+ proliferation from larval to mid-pupal stages, *DeltaRNAi* animals lose Imp expression early. Despite the loss of this early temporal factor, we do not see expression of its reciprocally inhibitory partner Syp. Furthermore, E93 expression is also delayed, raising the possibility that an extrinsic cue such as a pulse of the steroid hormone ecdysone is lacking so that the temporal factor progression is unable to proceed. We have shown that the Notch pathway positively regulates temporal patterning programs in a lineage specific manner as the effects seen in MBNBs are different than other CBNBs (Sood et al., 2023). At late timepoints when Notch signaling is perturbed, primarily through Dl knockdown, we also see a large amount of other CBNBs persisting past their typical termination time point (Fig. 1F). This highlights differences between NB lineages in the central brain.

### Apoptosis accounts for premature MBNB elimination

Due to shifts in temporal patterning primarily through extinction of imp expression prematurely, MBNBs are cycling through their lifecycle faster leading to early elimination. We assessed how MBNBs may be undergoing premature termination by blocking the apoptotic cell death pathway with a synthetic transgene against pro-apoptotic genes reaper, hid, and grim. We found that MBNBs were able to persist into adulthood when death was blocked in a *DeltaRNAi* animal and showed expression of the proliferative marker PCNA:GFP. We also assessed how the premature loss of the MBNBs affects the important MB structure by staining the ⍺, β, and γ neurons. We found that without Delta expression, we lose the late born ⍺ and β progeny resulting in a malformed MB. However, when we again block death by combining *UAS-miRHG* with *DlRNAi* we see more of this later born progeny and a more correctly formed γ lobe than in *DlRNAi* brains alone.

### Eyeless contributes to lineage specific differences in Notch activity

Finally, we assessed what lineage specific factor could cause such different phenotypes of premature elimination in MBNBs and ectopic persistence in other CBNBs. The evolutionarily conserved gene, Ey is specific to MBNBs, and we have previously observed phenotypes of premature loss under *EyRNAi* conditions (unpublished). This finding provides an excellent opportunity for future research into the role of Ey in regulating Notch activity in MBNBs, and how lineage specific differences between CBNBs arise.

Together, this has led us to conclude that the Notch pathway is important for proper MBNB proliferation and resulting MB morphology. Notch and its ligand Delta interact with the known temporal program of MBNBs to regulate when each type of progeny is produced and in the correct volume. This work also offers an opportunity to understand how other well-known signaling pathways may play a role in regulation of neurodevelopment through downstream temporal patterning genes. This is a novel component of MBNB temporal regulation that has not yet been recognized and allows for a deeper understanding of the genes involved in intrinsic control of neuroblast proliferation.

## Methods

### Fly Stocks

Fly stocks utilized, and their source are listed in Supplemental Table 1.

### Animal Husbandry

All animals were reared in uncrowded conditions at 25°C. Animals were staged from freshly hatched for larval time points and white pre-pupae for pupal time points.

### Clonal Induction

Frt-flp and MARCM clones were heat shocked at 37°C for 10-30 minutes during the first larval instar after staging at freshly hatched.

### Immunofluorescence and Confocal Imaging

Larval, pupal, and adult brains were dissected according to Pahl et al., 2019. Dissected brains were fixed in a solution of 4% paraformaldehyde in PEM buffer for 20 mins (larvae) or 30 mins (pupae), followed by a series of washes in PBST (1X PBS + 0.1% Triton X-100). Blocking solution of 10% normal goat serum (NGS) in PBST was applied and tissues were stored at 4°C overnight. Primary antibodies were applied and washed followed by Alexa-Fluor conjugated secondary antibodies (Supplemental Table 2). Secondary antibodies were washed, and brains were placed in anti-fade glycerol solution overnight prior to imaging. Z-stacks were taken of each central brain hemisphere by a Leica SP8 laser scanning confocal microscope using a 63x/1.4 oil immersion objective.

### Software and Data Analysis

Images were analyzed using Fiji ImageJ and processed using Adobe Photoshop. Figures were assembled using Adobe Illustrator. MBNBs were identified by nuclear dpn staining and superficial location along the dorsal surface of the brain. Cell size was calculated using Image J’s line tool to draw a cross hair across the NB cell body and the average of those two values was recorded in Graphpad Prism. Sample sizes are listed within the bars of all charts. All data is represented as mean ± standard error of the mean and statistical significance was determined using unpaired two-tailed student’s t-tests or ANOVAs in Graphpad Prism 9.

## Acknowledgements

We would like to thank the members of the Siegrist Lab for their help and feedback on this project.

## Figure Legends

**Supplementary Figure 1.**
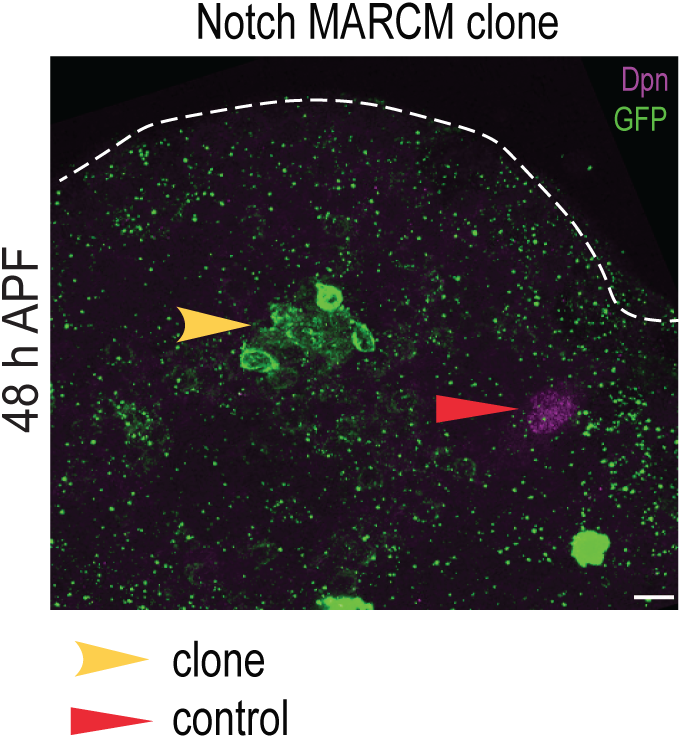
Notch MARCM clones show MBNB premature elimination phenotype. Maximum intensity projections of MARCM Clones at 48 h APF show a premature loss of Dpn+ MBNBs while control MBNBs remain Dpn+. GFP represents the clonal lineage from the eliminated MBNB. Scale bar = 10 μm.

## Abbreviations

ALH: after larval hatching
APF: after pupal formation
Br: broad
BTB-ZF: broad complex, tramtrack, bric-a-brac zinc finger
Cas: castor
CBNB: central brain neuroblast
Chinmo: chronologically inappropriate morphogenesis
CNS: central nervous system
Cx: calyx
Dl: delta
Dpn: deadpan
E93: ecdysone induced protein 93
E(spl): enhancer of split
Ey: Eyeless
FasII: fasciclin II
FE: freshly eclosed
GMC: ganglion mother cell\
Grh: grainyhead
Hb: hunchback
Imp: insulin-like growth factor II
KC: kenyon cell
Kr: kruppel
Mamo: maternal gene required for meiosis
MARCM: mosaic analysis with a repressible cell marker
MB: mushroom body
MBNB: mushroom body neuroblast
N: notch
NB: neuroblast
NGS: normal goat serum
NICD: notch intracellular domain
NSC: neural stem cell
PBS-T: phosphate buffered saline triton
PCNA:GFP: proliferating cell nuclear antigen green fluorescent protein
Pd: pedunculus
PFA: paraformaldehyde
PI3K: phosphatidylinositol 3 kinase
RHG: reaper, hid, grim
Scrib: scribble
Syp: syncrip

---

**Table 1.**

| Figure | Genotype |
| --- | --- |
| 1A, 1D, 4A-B<br>Control | *WornGAL4/+ (Oregon R); pcnaGFP/+ (Oregon R)* |
| 1B, 1E | *WornGAL4/+; pcnaGFP/UAS-NotchRNAi* |
| 1C, 1F, 3A | *worGAL4/+; pcnaGFP/UAS-DeltaRNAi* |
| 2A-B | *hsFlp; repoGAL80; UAS-DeltaRNAi/Act5c-FRT-CD2-FRT-Gal4, UAS-GFP* |
| 2C | *hsFlp; repoGAL80; UAS-DeltaRNAi/Act5c-FRT-CD2-FRT-Gal4, UAS-RFP* |
| 3B | *WorGal4/+; UAS-DeltaRNAi; UASmiRHG* |
| 4E-F | *WornGAL4/+; pcnaGFP/UAS-DeltaRNAi; UASmiRHG/+* |
| 4C-D | *worGAL4,tubGAL80(ts)/+; pcnaGFP/UAS-DeltaRNAi* |
| 5A-C | *hsflp; UAS-EyRNAi; E(spl)GFP/ Act5c-FRT-CD2-FRT-Gal4, UAS-RFP* |
| Supp. 1 | *hsflp, tubgal80, FRT19A/Notch55e11 FRT19A ; tubGal4, UASmCD8GFP/+* |

## KEY RESOURCES TABLE

**KEY RESOURCES TABLE:**
| Reagent | Source | Identifier |
| --- | --- | --- |
| **ANTIBODIES** |  |  |
| rat ant-Dpn (1:100) | Abcam | ab195173 |
| chicken anti-GFP (1:500) | Abcam | ab13970 |
| rabbit anti-Scribble (1:500) | Gift from Chris Q. Doe |  |
| guinea pig anti-E93 (1:250) | Gift from Chris Q. Doe |  |
| rabbit anti-Imp (1:250) | Gift from Paul MacDonald |  |
| rat anti-Imp (1:250) | Gift from Claude Desplan |  |
| goat anti-chicken Alexa 488 (1:300) | Thermo Fisher Scientific | A32931 |
| goat anti-rat Alexa 555 (1:300) | Thermo Fisher Scientific | A48263 |
| goat anti-rat Alexa 647 (1:300) | Thermo Fisher Scientific | A48265 |
| goat anti-rabbit Alexa 405 (1:300) | Thermo Fisher Scientific | A48254 |
| goat anti-rabbit Alexa 555 (1:300) | Thermo Fisher Scientific | A21428 |
| goat anti-rabbit Alexa 633 (1:300) | Thermo Fisher Scientific | A21071 |
| goat anti-guinea pig Alexa 488 (1:300) | Thermo Fisher Scientific | A11073 |
| goat anti-guinea pig Alexa 555 (1:300) | Thermo Fisher Scientific | A21435 |
| **CHEMICALS** |  |  |
| SlowFade™ Diamond antifade reagent | Invitrogen | Catalog # S36963 |
| SlowFade™ Gold antifade reagent | Invitrogen | Catalog # S36937 |
| Normal Goat Serum | Thermo Fisher Scientific | Catalog # 31873 |
| Paraformaldehyde 16% solution EM grade | Electron Microscopy Sciences | Catalog # 15710 |
| Triton X-100 | Sigma | Catalog # T9284 |
| **SOFTWARE** |  |  |
| ImageJ/Fiji | Fiji | http://fiji.sc/ |
| LAS X | Leica Microsystems | https://www.leica-microsystems.com/products/microscope-software/details/product/leica-las-x-ls/ |
| Prism 9 | Graphpad | https://www.graphpad.com/scientific-software/prism/ |
| Photoshop 2022 | Adobe | https://www.adobe.com/products/photoshop.html |
| Illustrator 2022 | Adobe | https://www.adobe.com/products/illustrator.html |

| **EXPERIMENTAL MODEL: *Drosophila melanogaster*** |  |  |
| --- | --- | --- |
| Oregon R | Bloomington Drosophila Stock Center | 5 |
| *wor-Gal4* | (Albertson and Doe, 2003) |  |
| *tubulin-Gal80(ts)* | Bloomington Drosophila Stock Center | 7108 |
| *UAS-Notch RNAi (HMS00001)* | Bloomington Drosophila Stock Center | 33611 |
| *UAS-Delta RNAi (HMS01309)* | Bloomington Drosophila Stock Center | 34322 |
| *pcna-GFP* | (Thacker et al., 2003) |  |
| *Act5c-FRT-CD2-FRT-Gal4, UAS-RFP* | Bloomington Drosophila Stock Center | 30558 |
| *Act5c-FRT-CD2-FRT-Gal4, UAS-GFP* | gift from Iswar Hariharan |  |

